# Comparative genomic evidence for self-domestication in *Homo sapiens*

**DOI:** 10.1101/125799

**Authors:** Constantina Theofanopoulou, Simone Gastaldon, Thomas O’Rourke, Bridget D. Samuels, Angela Messner, Pedro Tiago Martins, Francesco Delogu, Saleh Alamri, Boeckx Cedric

## Abstract

This study identifies and analyzes statistically significant overlaps between selective sweep screens in anatomically modern humans and several domesticated species. The results obtained suggest that (paleo-) genomic data can be exploited to complement the fossil record and support the idea of self-domestication in *Homo sapiens,* a process that likely intensified as our species populated its niche. Our analysis lends support to attempts to capture the “domestication syndrome” in terms of alterations to certain signaling pathways and cell lineages, such as the neural crest.

## Introduction

Recent advances in genomics, coupled with other sources of information, offer new opportunities to test long-standing hypotheses about human evolution. Especially in the domain of cognition, the retrieval of ancient DNA could, with the help of well-articulated linking hypotheses connecting genes, brain and cognition, shed light on the emergence of ‘cognitive modernity’. It is in this context that we would like to adduce evidence for an old hypothesis about the evolution of our species: self-domestication. As is well-documented^1^, several scholars have entertained the idea that anatomically modern humans (AMH) were self-domesticated. More recently, Hare^2^ articulated a more solid hypothesis bringing together pieces of strongly suggestive evidence.

It is clear that our species is unusually dependent on its socio-cultural environment to prosper. As Tomasello has stressed on numerous occasions (e.g.,^3^), our instinct to cooperate, possibly facilitated by a decrease in emotional reactivity^4^, marks us as ‘special’, certainly among primates, with the possible exception of bonobos, which have been claimed to be ‘self-domesticated’^5^. Our social tolerance and instinct to learn from members of our community are perhaps most obvious in the domain of language acquisition, which has led to claims that a self-domestication process in our species may have gone hand in hand with the emergence of a fully modern ‘language-ready’ brain^6–8^, and the construction and functioning of our cultural niche^1,9^. Apart from this behavioral uniqueness, several anatomical characteristics, especially when contrasted with what we know about Neanderthals (the best-studied example of archaic *Homo*), are reminiscent of what one finds in comparisons of domesticated species and their wild counterparts.

In order to detect the effects of domestication, data must come from comparisons of a domesticated species with “either their direct wild-living ancestor or close relatives if the ancestor is no longer extant”^1^. In the case of AMH, since there is no wild extant counterpart available, the obvious comparanda include our closest living relatives, i.e., the great apes, and extinct species of the genus *Homo* to the extent that relevant data can be extracted from the fossil record. Crucially for the purposes of this paper, we now have high-quality genomes for our closest extinct relatives, allowing for a more focused comparison. These comparisons can shed light on when the self-domestication event was initiated in our human lineage. We contend (*contra*^10^) that self-domestication coincided with the emergence of AMH (*sensu*^11^: specimens sharing a significant number of derived features in the skeleton with extant members of our species), because, as will be discussed below, some of the critical traits associated with the domestication process are defining features of our species’ anatomical modernity, although of course it is very likely the case that this self-domestication process intensified as our species expanded geographically and demographically.

Domesticated species display a range of anatomical and behavioral characteristics that set them apart from their wild counterparts: depigmentation; floppy, reduced ears; shorter muzzles; curly tails; smaller teeth; smaller cranial capacity (and concomitant brain size reduction); neotenous (juvenile) behavior; reduction of sexual dimorphism (feminization); docility; and more frequent estrous cycles. Of course, not all of these characteristics are found in all domesticates, but many of them are indeed present to some extent in all domesticates^12^. This constellation of features is known as the “domestication syndrome”, and has been hypothesized to arise from a mild deficit of neural crest cells^13^, to which we return below.

Many of the anatomical changes associated with domestication describe some of the well-known anatomical differences between AMH and Neanderthals (see Figure 1). The two species display different ontogenetic trajectories^14,15^ resulting in craniofacial differences that invariably lead to a more ‘gracile’, ‘juvenile’ profile in AMH relative to Neanderthals. It is well-established that prognathism is significantly reduced in our species^15,16^. Brow ridges and nasal projections are smaller in AMH than in our most closely related (extinct) relatives^10^, as is our cranial capacity^17^ and our tooth size^18,19^. This profile is sometimes called ‘feminized’^10^, and is associated with an overall reduction of sexual dimorphism, which is a trait associated with domestication^5^. The process of ‘feminization’ (reduction of androgen levels and rise in estrogen levels^10^) is often associated with reduced reactivity of the hypothalamus-pituitary-adrenal axis^20^, a physiological trait thought to be critical for domestication^13,21^. Evidence from digit ratio comparisons—a measure of prenatal androgen exposure, with lower second digit:fourth digit ratios pointing to higher prenatal androgen exposure, and therefore potentially a higher proclivity for aggressiveness^22^—further suggests that Neanderthals had higher prenatal androgen exposure than AMH^23^. Additional differences in other traits associated with domestication may exist, but there are either obvious confounding factors (e.g., geography for pigmentation) involved, or the data are subject to more controversial interpretation (e.g., in the case of reproductive cycle changes^24^) than the features reviewed above.

**Figure 1.**
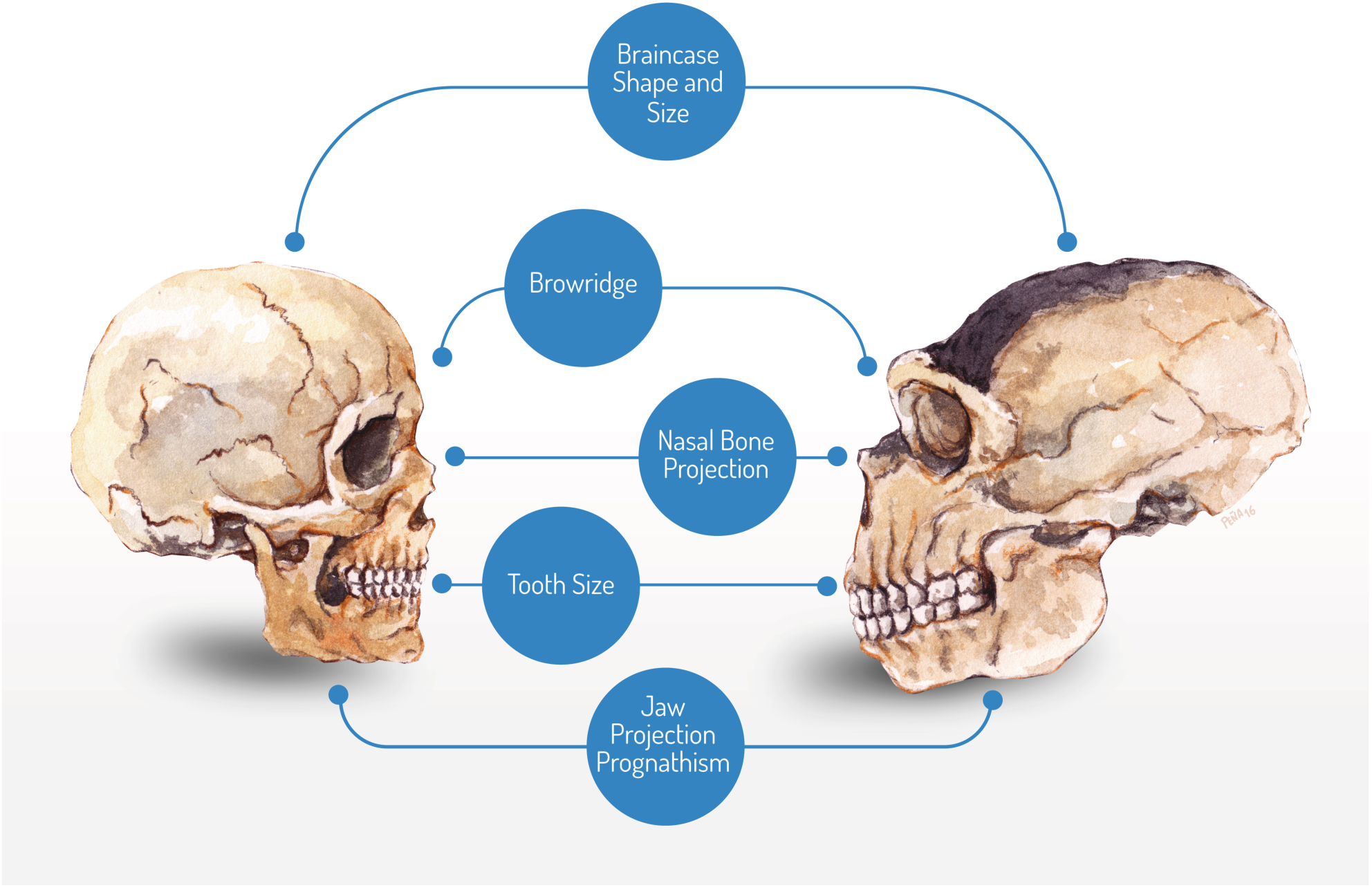
Salient craniofacial differences between Neanderthal and AMH, suggestive of domestication in the latter

The central claim of this paper is that there is an additional layer of evidence in favor of self-domestication in anatomically modern humans that can be constructed from findings from (paleo-)genomics. We now have detailed information concerning the genomes of several domesticated species and their wild counterparts^25^, as well as high-quality data from archaic human genomes^26^. This information offers the opportunity to test for the existence of overlapping regions of putative signatures of selection associated with (self-)domestication. If found to exist, such overlaps would complement the anatomical data briefly reviewed above and suggest that the self-domestication hypothesis is a strong contender to account for key aspects of modern human cognition.

## Results

We examined the overlap of gene sets independently claimed to be under positive selection in AMH (when compared with Neanderthal/Denisovan) and several domesticates for which detailed genetic information is available: dog (*Canis familiaris*), cat (*Felis catus*), horse (*Equus caballus*) and taurine cattle (*Bos taurus*). The pool of domesticates chosen yielded a total of 691 genes, and the total AMH pool, 742 genes. The intersection size (i.e., the number of genes that are positively selected or under selective sweep both in AMH and in one or more domesticate) returns a statistically significant number of 41 (*p* < 0.01) (See Table S2 and Figure 2). We confirmed the significance of this result with a Monte Carlo simulation of 1,000,000 trials, in which samples of 691 and 742 genes were randomly selected (with no replacement) from a pool of 19,500. The simulation confirmed that an intersection size greater than or equal to 41 is highly significant (*p* = 0.0003).

**Figure 2.**
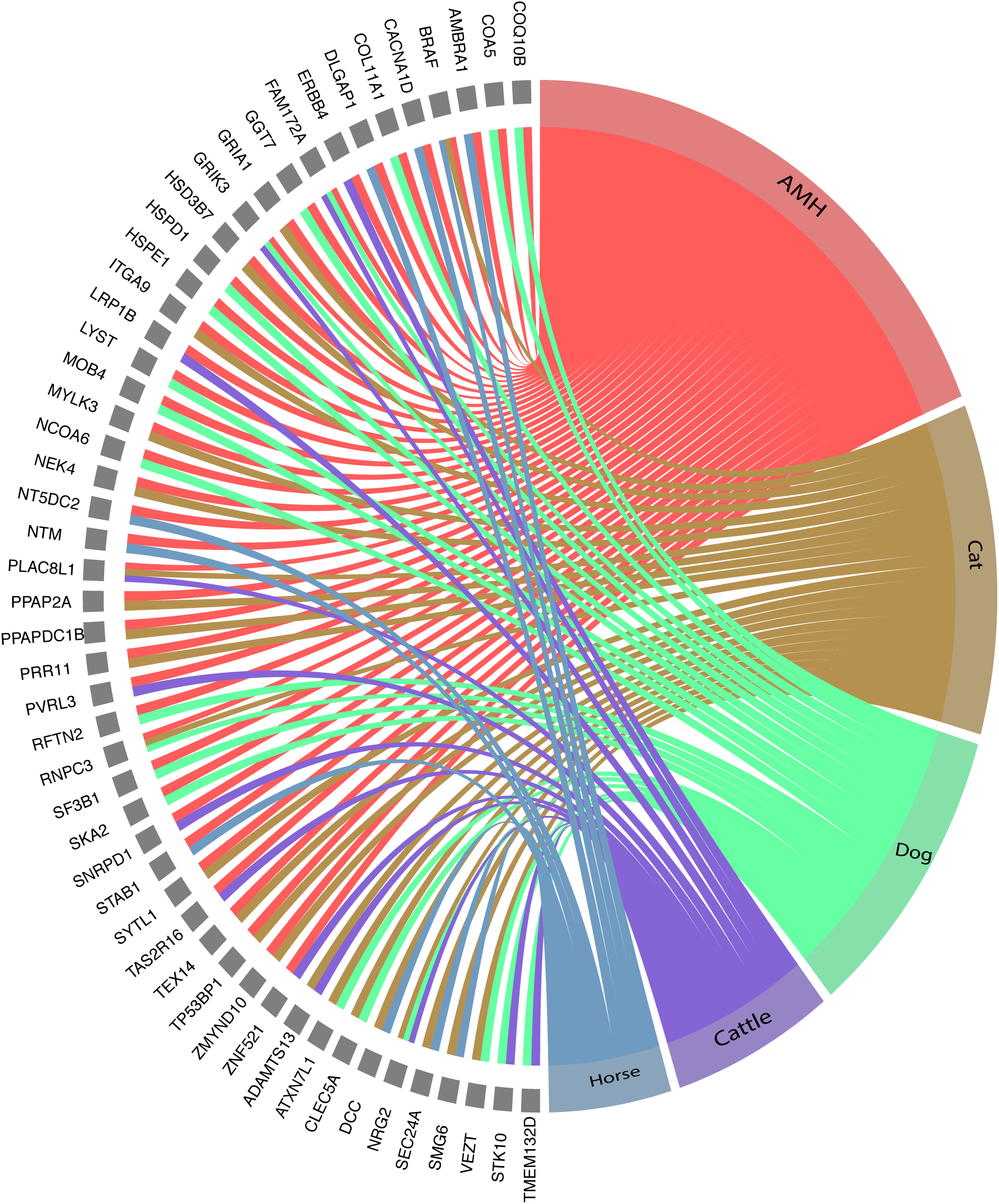
Graphical representation of the overlap identified among gene lists discussed in the text

This situation contrasted with the modest (statistically insignificant) overlaps between the domesticates and several Great Apes for which selective sweep screens were available: chimpanzee (*Pan t. troglodytes*), orangutan (*Pongo abelii*), and gorilla (*G. g. gorilla*) (Table S4).

Intersections between domesticates (15 genes in total, see Table S1) were tested, revealing a significant overlap between genes selected in the dog and in cattle (*p* < 0.01). Furthermore, tests between AMH and each domesticate showed significant overlaps with the dog (*v* = 15, *p* < 0. 05) and with cattle (*v* = 9, *p* < 0. 01). Since we pooled data for positive selection and selective sweeps in AMH from different sources, intersection tests were carried out between the domestication pool and the pool of each AMH dataset used in this study. A significant intersection was found with the data in^27^ (*p* < 0.05) and with the combined data from^27^ and^28^ (*p* < 0.05). Intersections between domesticates were also tested, revealing a significant overlap between genes selected in the dog and in cattle (*p* < 0.01).

Though no gene was found to be shared across all domesticated species studied here as well as AMH, this is not necessarily expected—domestication is known to proceed through various routes (see Discussion), and is thus not a uniform affair. However, common pathways can be identified, as can genes that may have contributed to domestication events that we think deserve special attention. Five genes were found to be associated with signals of positive selection in AMH and multiple domesticated species (Table S3): *RNPC3, FAM172A, PLAC8L1, GRIK3* and *BRAF*.

*RNPC3* shows evidence of positive selection in the dog, cat, and AMH. *RNPC3* is one of only two genes with more than one putatively causal variant fixed between dogs and wolves (the other is a gene of unknown function)^29^. Mutations in *RNPC3* cause growth hormone deficiencies in humans resulting from pituitary hypoplasia^30,31^. In a similar vein, a gene showing an AMH-specific amino acid change and associated with a strong positive selection signal in AMH and in dogs, *NCOA6*, is a nuclear receptor coactivator that directly binds nuclear receptors and stimulates the transcriptional activities in a hormone-dependent fashion.

*FAM172A*, selected for in dogs, cattle, and AMH, may perhaps be worthy of note given its position on chromosome 5 neighboring *NR2F1*, which plays a role in regulating neural crest specifier genes and has undergone selection in AMHs^32,33^; The functionally related nuclear receptor *NR2F2*, involved in regulating embryonic stem-cell differentiation^34^, and implicated in neural crest development, has been selected for in the domesticated fox^35^.

*PLAC8L1* is selected for in more than one domesticate species (cat, cattle) as well as in AMH, but there is only sparse evidence concerning its function. An autistic patient has been noted as having a microdeletion at chromosome 5q32, a location which includes *PLAC8L1^36^*.

The two remaining genes in Table S3, *BRAF* and *GRIK3*, deserve special attention.

### ERK pathway

*BRAF*, selected for in the cat, horse, and AMH, is an important member of the ERK/MAPK signaling pathway, which has been shown to play a key role in synaptic plasticity, memory, and learning^37^, and which, when disrupted, can lead to a broad range of syndromes comprising craniofacial defects and cognitive deficits^38^. *BRAF* is upstream of *ERK2*, which plays a critical role in neural crest development^39^ and regulates neuronal gene expression in both the neocortex and hippocampus^37^. Both *BRAF* and *ERK2* inactivation can bring about syndromic symptoms by disrupting neural crest development^39^. *BRAF* is implicated in Noonan, Leopard, and Cardiofaciocutaneous syndromes, typical symptoms of which include prominent forehead, bitemporal narrowing, hypertelorism, short stature, among other skeletal, cardiac, and craniofacial anomalies, frequently accompanied by moderate to severe mental retardation^40,41^. *BRAF*, interacts with other domestication-related genes, including *YWHAH* (dog), *PPP2CA* (a neural crest-related gene, selected for in the horse), and *HER4/ERBB4*, another neural crest gene selected for in cattle and AMHs. Upstream of *BRAF, SOS1*, selected in domesticated foxes, affects MAPK signaling, bringing about Noonan phenotypes^42^. Noonan-syndrome-like phenotypes are associated with several genes that appear to have undergone selective sweeps in AMH. For instance, *CBL* is located in a region showing signal of a strong selective sweep in AMH compared to Altai Neanderthals^27^, and, when mutated, has been shown to give rise to a Noonan syndrome-like disorder^43^.

As mentioned above, the AMH-selected neuregulin (NRG) receptor *ERBB4* is part of the ERK/MAPK pathway and negatively regulates ERK via upstream phosphorylation of Raf-1^44^. Loss of function of *Erbb4* in mice has been shown to cause defects in hindbrain cranial neural crest cell pathfinding, including a caudal elongation of the trigeminal and geniculate ganglia^45^. This suggests a plausible role for *ERBB4* in preventing caudal extension in the derived AMH skull. *ERBB4* is one of many neural crest-related genes associated with selective signals in AMH (e.g., *SNAI2*^27^, *CITED2*^27^, *PRDM10*^46,47^; and other genes discussed in^6^), some of which show fixed or nearly fixed amino acid changes compared to Neanderthals. In addition, *NRG2* was the only gene that was found to be selected in three of the four domesticated species in our study: cat, cattle, and dog. NRG4 was selected in cattle, and NRG3, in AMH. Incidentally, *NRG3* copy number and single nucleotide variants have been associated with Hirschsprung disease^48,49^. This disease is very relevant in the context of domestication, as it affects the neural crest, associated with domestication syndrome^6,13^. Quite a few genes associated with selective sweeps in AMH examined here (among them, *RET, ZEB2*, and *SLIT2*) have been linked to the disease^50,51^.

Enhanced ERBB4 signaling has been implicated in Angelman syndrome, an autism spectrum disorder marked by behavioral traits such as increased desire for social interaction, developmental delay, severe speech impairment, and a happy demeanour, although aggressive behavior has sometimes been reported^52–54^. Angelman-syndrome-like phenotypes are frequently associated with genes investigated here. One such observation concerns “the most intriguing variant fixed between dogs and wolves”^29^, which is found in the 3’-UTR of *SLC9A6*. This gene encodes sodium/hydrogen exchanger protein 6, which is part of a network related to the plasticity of glutaminergic neurons^55^.

Cagan and Blass^29^ note that loss-of-function mutations in this gene in humans can lead to Christianson syndrome, also known as “Angelman-like syndrome”. Phenotypes typical of these patients include cognitive developmental delays, absence of speech, stereotyped repetitive hand movements, and postnatal microcephaly with a narrow face. Christianson syndrome is frequently characterized by a happy disposition with easily provoked laughter and smiling, an open mouth with excessive drooling and frequent visual fixation on hands. Several of these phenotypes resemble those that distinguish dogs from wolves.

It is noteworthy that in pathway analyses carried out on the lists of genes in Tables S1 and S2, as well as the list of genes with amino acid replacement substitutions fixed in AMHs and absent in archaic humans^56^, ERKs are among the most significant downstream targets of the interacting selected genes in each.

As a final note on the ERK pathway, we highlight the presence of *CACNA1D* in Table S2. *CACNA1D* is a neural cell adhesion molecule that contributes to cell migration via activation of MAPK/ERK signaling^57^. This gene is one of several axon-guidance molecules we identified in our study. It is highly expressed in the adrenal glands^58^, and, when mutated, gives rise to cerebral palsy/motor disorders^59^. It has been linked to auditory processing^60^, and said to be among the positively selected genes in some vocal learners^61^.

### Glutamate receptors

The glutamate receptor GRIK3 occurs in the AMH, dog, and cattle lists. interacts with other glutamate receptors selected for in the horse (*GRID1*) and cat (*GRIA1/2*). Polymorphisms in *GRIK3* and *GRID1* have been implicated in schizophrenia^62,63^, and *GRID1* neighbors *NRG3* (discussed above) at the schizophrenia susceptibility loci 10q22-q23^64^. Developmental delays and craniofacial anomalies associated with a loss of genetic material at the *NRG3* locus accompanied by a gain of material at the *DLGAP1* site have also been reported^65^. *DLGAP1*, a scaffold-protein-coding gene at the postsynaptic density, is in fact selected for in AMHs and in the horse. This gene has been implicated in obsessive-compulsive disorders and interacts significantly with Shank proteins, mutations in which have been linked to autism spectrum disorders with impaired social interaction and communication^66–68^. DLGAP1 interacts with the glutamate receptor GRIK2, implicated in obsessive-compulsive disorders^69^.

Previous work^70^ already pointed out that genes involved in glutamate metabolism show the greatest population differentiation by whole-genome comparison of dogs and wolves. Although such changes may be implicated in fear response differences between the dog and the wolf populations, Li et al. argue for a role in increasing excitatory synaptic plasticity in dogs rather than reducing fear response. As they point out, changes related to synaptic plasticity may have a significant impact on learning and memory. This is certainly true for cognitive specializations in humans, like language, since glutamate receptors have been shown to be differentially regulated in brain regions associated with vocal learning^71^.

### Genes selected for in multiple domesticates but not AMH

It is worth considering those genes selected for across domesticates, independently of their selection in AMH, for different reasons. First, our aim here is to explore the extent to which self-domesticating processes in humans may have contributed to our species’ anatomical, cognitive, and behavioral make-up. Uncovering strong candidate genes for (albeit often different) domesticating processes in other well-studied domesticated species is a promising way to pursue this goal. The strongest of these candidate genes should be those that have been selected for across different species. Those genes selected for only in certain domesticates, but which strongly interact with genes selected across domesticated species, may prove central to a relevant domesticating process, given that these interactions may shed special light on relevant phenotypic traits. Similarly, certain genes selected for across different domesticates may have trong interactions with other genes that *are* selected for in AMHs. We wish to highlight some of these here (for a full list, see Table S1).

*DCC (DCC Netrin 1 receptor)*, an axon-guidance mediator and neural crest-related gene, selected for in both the horse and the cat, interacts strongly with *DSCAM (Down Syndrome Cell Adhesion Molecule*), another axon-guidance and neural crest-related gene, selected for in cattle. Of key significance is interaction of *DCC* with the Slit/Robo pathway, especially given the latter’s proposed involvement in vocal learning^72,73^ and the selection of both *ROBO2* and *SLIT2* in AMHs^47^. (*ROBO1* is also selected for in cattle.) ROBO silences the effect of Netrin 1 to DCC, allowing SLIT2 to bind to this ligand and enabling axon pathfinding in the developing brain^74,75^. *DCC* is involved in the organization of dopaminergic circuits within the cortex^76^, and several association studies have identified *DCC* as a promising candidate for schizophrenia^77^. Importantly, an AMH-specific hCONDEL exists in a region upstream of *DCC*, although it is shared with Neanderthals^78^. However, a detailed examination on this gene on both the modern and archaic lines, reveals an accummulation of changes on this gene in AMH. In addition, several genes showing AMH-specific amino-acid substitutions, such as *NOVA1* and *RASA1*, both involved in neuronal development, are known to interact with *DCC*^79,80^, and could regulate it in a species-specific fashion. *RASA1* is associated with a strong selective signal in AMH, and has been shown to mediate Netrin 1-induced cortical axon outgrowth and guidance^80^. Together with the glutamate receptor changes discussed above, such modifications may have played an important role in generating aspects of the cognitive profile associated with modern humans, including a full-fledged language-ready brain.

We found several collagen-type genes selected for across domesticates. *COL22A1*, a gene selected for in the horse, significantly interacts with various similar genes selected for in other domesticates, particularly in the cat, including *COL11A1*, selected for in dogs and AMHs. *COL22A1* and *COL11A1* exhibit increased expression in the bone tissue and hippocampus of mice with some of the symptoms of Kleefstra Syndrome (developmental delay, hypotonia, and craniofacial abnormalities), which are often accompanied by autistic symptoms and intellectual disability in humans^81,82^.

### Archaic-derived alleles

To the best of our knowledge, no comprehensive selective sweep analysis exists for neanderthals. We examined the genes associated with archaic-derived alleles^83^ and found no genes in Table S1 display reported archaic-derived alleles. While this could be due to the smaller number of archaic-specific SNCs known at the time of writing, we find this to be an important contrast with the situation that obtains with AMH, in light of the self-domestication hypothesis. It is striking that Castellano et al.^83^ highlight genes involved in skeletal development and associated with aggressive phenotypes.

We have been able to find data concerning nearly fixed ancestral or derived SNPs in archaic lineages that crop up as variants in modern-day populations. Despite the many confounding factors as to how the relevant mutated genes might interact in different genetic contexts, one might still expect certain archaic-selected SNPs to exhibit somewhat ‘underdomesticated’ phenotypes when occurring as AMH variants. In this sense, mutations imitating ancestral SNPs found in archaic lineages may be able to tell us a great deal about the evolution of our lineage, by allowing us to glimpse some aspects of the ancestral genotype. Among those mutations we found, there is an ancestral S330A mutation of *SLITRK1* that may be involved in obsessive-compulsive disorders like Tourette’s Syndrome^84,85^. Different amino acid changes around the site of an AMH-specific derived protein ADSL (A429V) can bring about adenylosuccinate lyase deficiency (R426H; D430N^86^), the symptoms of which include developmental delay, autistic-like traits, aggressiveness, and microcephaly^87^.

## Discussion

As several scholars have pointed out^5,12,88^, there are several routes to domestication. These include the directed pathway (human-assisted amplification of some desired trait in a species), the prey pathway (animals previously hunted in the wild, and subsequently managed in herds), and the commensal pathway (in which animals are drawn to humans by attractive food sources, and over time domesticate themselves). As a result, we should expect genes targeted by a domestication process to differ considerably across species. Nevertheless, reviewing the molecular events associated with domestication reveals common themes, with significant numbers of genes related to brain function and behavior, anatomy, and diet, across domesticates. This is consistent with the view that domestication may be best represented as a spectrum or continuum^89^, with a polygenic basis and non-uniform symptomology. This state of affairs is well reflected in the study conducted by Albert et al. (2012)^90^, who found significant brain expression changes across domesticates, with the majority of these changes being species-specific.

Because of these findings, we find the overlaps listed in Tables S1-S2 and the associated functions and pathways discussed in the Results section all the more relevant, especially because they converge to a large extent with what is to be expected from the neural-crest-based hypothesis^13^ put forth to capture the common mechanistic basis of domestication events. According to Wilkins et al. (2014)^13^, a disruption in neural crest developmental programs might be the source of changes spanning multiple organ systems and morphological structures, and the genes examined here seem to broadly support this view. It is quite possible that a neural crest-based explanation won’t apply to all domesticates, as suggested in^12^, but we find it significant that this hypothesis finds its strongest support in domesticated species like dogs, which have been argued to be self-domesticated^2^. Recall that the goal of the present study was not to provide molecular evidence for a general theory of domestication, but rather to identify domestication-related pathways that could be suggestive of a self-domestication process in AMH. The fact that we find neural-crest-related changed in AMH compared to Neanderthals/Denisovans, and that such changes are also found in another species hypothesized to have undergone a self-domestication process, reinforces our hypothesis that self-domestication took place in our species.

Apart from neural crest-related genes or pathways, we identified common themes pertaining to neuronal development, synaptic plasticity and enhanced learning (categories often mentioned in independent studies on selective sweeps in AMH; e.g.,^47^). These results are in line with claims in other studies on domestication^91–94^, where categories like ‘neurological process’ frequently stand out strongly in gene ontology category enrichment analyses. For scientists interested in the evolution of human cognition, this is of great relevance. It lends credence to claims pairing domestication and a certain type of intelligence^95^, and it is not unreasonable to suspect that byproducts of the domestication process such as enhanced sensory-motor perceptual and learning pathways, may provide a foundation for more complex communicative abilities, including vocal learning abilities^7,96^.

In a similar vein, we want to point out that among the genes found in Tables S1-2, one finds multiple strong candidates for neurodevelopmental diseases and syndromes (as mentioned for several genes in the Results section). This could be seen as an additional piece of evidence suggestive of a self-domestication process in AMH. A build-up of deleterious alleles is documented across domesticated species when compared to their wild counterparts. Thus, there is a higher frequency of non-synonymous substitutions in the nuclear DNA of domesticated dogs relative to gray wolves^97^, and the same is true of their mitochondrial DNA^98^. A higher frequency of non-synonymous substitutions in domesticated yaks compared to the wild yaks has also been reported^99^. This build-up of deleterious alleles has been described as the ‘cost of domestication’^100^, which, if true, could be a byproduct of self-domestication in AMH, too. For instance, Buffil et al.^101^ points out that elevated aerobic metabolism, which may lead to the retention of ‘juvenile’ neuronal characteristics including elevated synaptic activity, may predispose these neurons to be more sensitive to oxidative stress and thus neurodegenerative diseases.

A study like the present one suffers from several limitations. While we have tried to make our comparisons as fair as possible, we have relied on genomic data that necessarily reflect the current state of the art for the various species we examined. The lists of genes associated with signals of positive selection are derived from the literature, and were generated using different analytical tools. While we have done our best to minimize the number of simplifying assumptions (see Methods), we must point out that even within a single species (e.g., AMH), no two studies completely agree on a definitive list. Indeed, in some cases, they produce lists of very different sizes. In addition, We may have missed important genes of interest due to the lack of information on them in the various databases we consulted. While it is to be hoped that some of these limitations will be overcome in the future, we think that the overlaps discussed in this study should encourage further detailed examination of these genes and the processes they take part in.

We could have been more strict about our notion of convergence, and restrict our attention to genes where the exact same difference (e.g., the same amino acid substitution) could be detected between species (for an early attempt along these lines, see^102^). But given that convergent evolution is often hypothesized to occur in the absence of this very strict notion of convergence — for instance, convergent evolution in the domain of vocal learning is related to non-identical changes in *FOXP2* across vocal learners^103^ — we feel justified in our approach.

As more data about selective sweeps become available, we think that it will be worth examining the extent to which our results carry over to other species like the bonobo, which has been claimed to be self-domesticated^5^. Likewise, more information about the molecular basis of the famous domestication experiment with the silver fox (*Vulpes vulpes)^104^* should prove valuable. As a matter of fact, the only study focusing on the genetic divergence between foxes that were selected for tame and aggressive behavior^35^ reveals an intriguing overlap between a set of SNPs significantly differentiating the two fox strains and among the genes noted in the silver fox data^35^ and a number of genes considered to fall within non-coding human accelerated regions (ncHARs)^105^. The most significant among these genes are those implicated in neural tube and forebrain development, and are listed by Racimo (2016)^28^ as having been selected from within a single region on chromosome 12 in the AMH Eurasian branch. The region containing all of these genes and no others has been deleted in at least one recorded adult patient with mental retardation and psychiatric problems^106^. Data of this kind have the potential to reinforce the argument presented in this study.

## Methods

### Data

To identify signatures of a self-domestication process in AMHs, we first constructed a list of genes associated with signs of positive selection in AMH compared to Neanderthals and Denisovans. We then compared this list to the genes independently argued to be associated with positive selection in domesticated species versus their wild counterparts, and examined the overlap between these two gene lists.

For AMH-Neanderthal/Denisovan comparisons, we made use of findings based on high-quality genome reconstructions, specifically: the list of genes in regions of putative selective sweeps, together with pathway and disease annotation in [27, Table S19b.1]; the list of genes from the top 20 candidate regions for the modern human ancestral branch in [28, Table 3]; and the extended list of genomic regions that are predicted to underlie positively selected human specific traits in [47, Supplemental File S1 and S2, Table 2, and Table S7].

We included in our study a range of domesticated species for which detailed genetic information is available. These species offer representative examples of the various routes to domestication^88^, as well as different temporal windows for domestication. The species include: dog (*Canis familiaris*)^29,93,107^, cat (*Felis catus)^108^*, horse (*Equus caballus)^109^*, and taurine cattle (*Bos taurus*)^110^. We homogenized the nomenclature across gene sets as best we could.

We also examined other species, including the rabbit (*Oryctolagus cuniculus*)^94^, and bonobo (*Pan paniscus*)^111^. In the end, the lists of selected genes for these species (compared to their wild counterparts) were too small to draw any firm conclusion. To help us understand domestication-related changes better, we made use of the comparison of two lines of rats (*Rattus norvegicus*) selected for tame and aggressive behaviour to identify genetic loci that differ between the lines^112^, the comparison of gene expression levels in the brains of domesticated and wild animals offered in^90^, genomic signatures of domestication in neurogenetic genes in *Drosophila melanogaster*, where neurogenetic genes have been claimed to be associated with signs of positive selection^91^, and the genetic divergence between foxes (*Vulpes vulpes*) that were selected for tame and aggressive behavior^35^.

For the Great Ape comparison — chimpanzee (*Pan t. troglodytes*), orangutan (*Pongo abelii*), and gorilla (*G. g. gorilla*) — we made use of the data in^113^ (positive and balancing selection and selective sweep data in Tables S6, S18(68), S19(69), S20(70), S24(74), and S97).

### Methods

In order to test the significance of the overlap between domestication-related genes and genes showing signals of positive selection and selective sweep in AMH, a Hypergeometric Intersection distribution test was performed by using the *R* software^114^ and the *R* package hint developed in^115^. A Hypergeometric Intersection distribution can be employed to compute the probability of picking an intersection of size *v* when drawing independently and without replacement from two sets *A* and *B* composed of objects of *n* categories, with *a* and *b* number of draws respectively (where *a ≠ b*)^115^.

As a model of our data we chose as a simplifying assumption *n* = 19,500 as the average number of protein-coding genes for all the species taken into consideration. From the original lists, we thus removed antisense RNA genes (non coding), miRNAs, and other non-coding transcripts/products listed in the original tables.

From this modeled genome, the domesticate pool (comprising cat, dog, cattle, and horse) draws a total of *a* = 691 genes, while the total AMH pool draws *b* = 742 genes. The resulting intersection size (i.e. the number of genes that are positively selected or under selective sweep both in AMH and in one or more domesticate) is *v* = 41. The `hint.test` function was then employed to test the significance of this intersection, obtaining *p* < 0.01.

A Monte Carlo simulation was performed using Matlab (MathWorks, Natick, MA) to confirm these results. Two random samples, of lengths 691 and 742, were drawn from a pool representing 19,500 genes using Matlab’s random number generation function. These simulated draws were performed 1,000,000 times and the percentage of trials in which the intersection was ≥41 was calculated. The results revealed that 0.33% of trials had intersections of this size.

Intersections between domesticates were tested, revealing a significant overlap between genes selected in dog and in cattle (*a* = 229, *b* = 78, *v* = 5, *p* < 0.01). Furthermore, tests between AMH and each domesticate showed significant overlaps with dog (*v* = 15, *p* < 0. 05) and with cattle (*v* = 9, *p* < 0. 01).

Since we pooled data for positive selection and selective sweep in AMH from different sources, intersection tests were carried out between the domestication pool and the pool of each AMH dataset used in this study. A significant intersection was found with the data in^27^ (*a* = 691, *b* = 108, *v* = 9; *p* < 0.05) and with the combined data from^27^ and^28^ (*a* = 691, *b* = 419, *v* = 24, *p* < 0.05).

Overlaps with domesticates were checked for Great Apes, using data from^113^. For chimpanzee (*Pan t. troglodytes), b* = 415 with *v* = 16; for orangutan (*Pongo abelii*), *b* = 500 with *v* = 20; for gorilla (*G. g. gorilla*), *b* = 426 with *v* = 12. The tests yielded non-significant results for all these intersections. Monte Carlo simulations, performed as described above, *mutatis mutandis*, showed that intersections of these sizes occurred in a large fraction of trials (40.11% of trials for chimpanzee; 32% for orangutan; 82.89% for gorilla). As in the case of AMH, overlaps with individual domesticates were tested, with no significant results.

We then examined the functions of the genes in Tables S1-3, paying close attention to the pathways in which they are involved, and to their interactions with other genes already highlighted in the domestication literature. In addition to performing an exhaustive PubMed (http://www.ncbi.nlm.nih.gov/pubmed) search on each of the genes, we drew on the information available in Genecards (http://genecards.org), Uniprot (http://www.uniprot.org/), String 10.0 (http://string-db.org), and Biogrid 3.4 (http://thebiogrid.org) to identify potential protein-protein interactions and Gene Ontology category enrichment signals. Additionally, we studied the networks and functional analyses generated by feeding our gene lists into QIAGEN’s Ingenuity Pathway Analysis software (https://www.qiagen.com/ingenuity).

Furthermore, we gathered information about the expression patterns of these genes, concentrating on those genes with relatively high expression in tissues such as brain, bone, and adrenal glands. For this, we relied on the following resources: Brainspan (http://www.brainspan.org), Human Brain Transcriptome (http://hbatlas.org), Bgee (http://bgee.org), Proteomics DB (https://proteomicsdb.org), Human Protein Atlas (http://www.proteinatlas.org), Gene Enrichment Profiler (http://xavierlab2.mgh.harvard.edu/EnrichmentProfiler/index.html), and GTex (http://www.gtexportal.org). For the information presented in the Supplementary Material, we consulted the following databases: KEGG Pathways and Disease (http://www.kegg.jp/kegg/), PANTHER (http://www.pantherdb.org), Reactome Pathway Database (http://www.reactome.org), OMIM (http://omim.org), and MalaCards (http://www.malacards.org/).

## Acknowledgements

We acknowledge the help of Nina Riddell in generating Figure 2. The drawing for Figure 1 was done by Francisco Peña. We thank Kay Sušelj for assistance with the Monte Carlo simulations.

## Author contributions statement

CB ad CTh formulated the initial hypothesis. SG and CB designed the present study. SG, FD, CB, PTM, and BDS selected the methods and performed the statistical analyses. Data collection: SG focused on genomic information related to the dog; PTM, to the cat; TOR, to cattle; AM, to the horse. CTh collected data on the bonobo and SA collected data on the rabbit, which were eventually not included in the present report. CB and CTh produced a first draft of the Introduction and Discussion; CB, TOR and SG produced a first draft of the Results section. CB, SG, and AM produced a first draft of the Methods section. CB and BDS thoroughly revised the drafts and produced the final version of the manuscript, with input from all authors. AM, SG, TOR, BDS, and CB prepared the supplementary material. CTh and SA prepared the illustrations. CB coordinated the study.

## Additional information

The authors declare no competing financial interests.

